# Effects of Ivermectin exposure on regeneration of *D. dorotocephala* planaria

**DOI:** 10.1101/186650

**Authors:** Nina N. Ferenc, Michael Levin

## Abstract

The ability of cells to communicate is essential during pattern formation, as they make decisions that drive growth and form. One mode of cellular signaling is via bioelectrical properties determined by the activity of ion channels. Several studies have shown a role for bioelectric signaling in planarian regeneration, but these have focused on *D. japonica* and *S. mediterranea.* It is not known how the alterations of ion channel activity would affect regeneration in other species of planaria. Here, we tested the effect of ivermectin (IVM), a chloride channel opener drug commonly used to combat heart worms, on regeneration in a new species of planaria: *D. dorotocephala.* Exposure to IVM during regeneration resulted in patterning abnormalities, such as bifurcated tails with partial heads, as well as delayed regeneration. By testing the effect of drugs that target resting potential on regenerative repair in novel model species, additional insight is gained on the comparative roles of ionic signaling across taxa.

## Introduction

Understanding how cells communicate toward making complex patterns is crucial to the development of novel regenerative therapies. Disturbances in this process, which manifest as birth defects, failures of regenerative repair, or cancer, provide important clues about the normal function of cellular decision-making in vivo. Biochemical and biophysical signals are utilized for patterning, and a number of model species facilitate the study of these processes [1–3].

Planaria have remarkable regenerative capabilities, allowing scientists to better understand the process in repairing a complex body via manipulation of patterning activity. These free-living flatworms serve as proof-of-principle that complex regeneration of body-parts after injury is possible throughout the lifespan [4, 5]. Thus, it is essential to understand the signals and biophysical properties that underlie patterning decisions in this model system. Planaria have a complex bodyplan with many types of organs and tissues, including a nervous system consisting of a brain connected by two nerve cords running along the body to the tail that are connected by transverse nerves [6]. After amputation, each section of the body activates cells called neoblasts - an adult stem cell population responsible for producing the cell types needed for regeneration [7]. Neoblasts are pluripotent, possessing the ability to give rise to various cell types [8]. These neoblasts proliferate and migrate to replenish the missing tissues by forming a blastema [6]. Planaria regenerate exactly what is missing and stop when the correct planarian target morphology has been achieved. Aside from specification of cell types from the planarian stem cells, regeneration requires the ability to place these cells in proper geometric relationship with each other - to produce appropriately located, sized, oriented, and patterned complex organs. This process is an active area of investigation, and a wide range of genetic and biophysical processes are involved [9–11]. In complement to the mainstream work on biochemical signaling in planarian regeneration, here we focused on the role of ionic controls of repair.

Like the neurons in the brain, cells in all tissues consist of electrical networks to process information and communicate with each other through the membrane in order to make critical patterning decisions, such as those required for regeneration [10]. Ion channels, which are regulated by numerous physiological signals (including resting potential itself), allow ions to cross the membrane creating voltage gradients across all tissues [10, 12].

Bioelectrical signaling via changes in cells’ plasma membrane voltage is an initiator of regeneration, and is a signaling modality that underlies complex patterning in numerous model species [10, 13, 14]. One method of altering membrane voltage for functional experiments is to open or close specific ion channels using drugs such as ivermectin (22,23-Dihydroxy-avermectin B). IVM binds to glutamate-gated chloride channels (GluCl) which increases the permeability of chloride ions across the cell membrane [15]; this reagent has been used to regulate bioelectric signaling in cancer [16], innervation from transplants [17], and brain patterning [18]. Therefore, exposure to IVM could result in alterations of normal repair processes, shedding light on the roles of chloride channels in regeneration. By altering pattern formation of the regenerating flatworms, analysis of anatomical outcomes will provide insights into the biophysical capacity of regeneration.

Malfunctioning ion channels have been linked to a range of birth defects (channelopathies) and cancer [2, 19–23], while modulation of resting potentials in vivo has been used to override default patterning outcomes in developmental, regenerative, and neoplastic contexts [26, 27]. New treatment approaches will require a better understanding of the endogenous function of such channels in order to control their contributions to the control of growth and morphology [24, 25]. Planarian species such as *D. japonica* and *S. mediterranea* have been recently used to reveal how ion transport and the resulting voltage gradients control pattern during regeneration [10, 16, 26–33]. However, differences have been reported in these two worm species’ phenotypes resulting from V_mem_ modulation, and it is not known how bioelectric signaling functions in other species of planaria. In order to have a more comprehensive understanding of bioelectrics across different species, we explored the role of the altered ion channel activity in the regenerative process using *D. dorotocephala.* These data for the first time reveal the dosage tolerance and regenerative outcomes of exposure to ivermectin in this species.

## Materials and Methods

### Animal husbandry

*D. dorotocephala* were supplied by Carolina Biological Supply Company and were starved ≥5 days before use to reduce metabolic variability amongst the worms [34]. Planaria were not fed for the duration of the experiment. Each concentration of planaria used 60 fragments of worms in order to collect adequate data and increase the statistical accuracy of results

### Preparation and pre-soak

Before receiving planaria, 15 petri dishes (60x15mm) were rinsed in water. Spring water (Deer Park) was used throughout, to avoid the toxicity that may be caused by the chlorine in tap water [35]. Then each dish was filled with 9.90mL of spring water using a Fisher Scientific 10mL sterile disposable serological pipets. The stock solution of 10mM of ivermectin in DMSO was diluted to 100μM to allow the pipettes to take in a more precise amount, thus avoiding possible errors [26]. DMSO was used as a solubility-enhancer of IVM in the aqueous solution [36, 37]. Using a VWR brand micropipette (100μL and 200μL) with the attached VWR’s Pipet tips, 100μL (1μM) of IVM was added to each dish except for the controls (0μM). The remaining IVM was separated equally into a total of four individual Eppendorf tubes to avoid thawing IVM aliquots multiple times. For the control (0μM), 1μM of DMSO (99.9% pure liquid Dimethyl sulfoxide by Nature’s Gift) was added to three petri dishes labeled “control” in order to standardize the concentration of all pre-soaks. All other concentrations received 1μM of IVM in DMSO into 9.90mL of water. The purpose of the soak exposure is to allow IVM to penetrate into the tissue of the planaria, so that the IVM will have an effect during the first phases of regeneration [26]. Upon arrival of planaria, 10 planaria were immediately transferred to each of the 15 petri dishes using a 3mL pipette without transferring any water. A cardboard box was used to cover the dishes as planaria are light-sensitive [38]. After 24 hours, all the water from every petri dish was removed using a 10mL pipette without extracting the planaria. Then the soak was refreshed by re-applying the soak measurements as before by adding water and IVM/DMSO to the designated dishes. After another 24 hours, all liquids from the dishes were removed. Another separate set of 15 petri dishes were rinsed with spring water and labeled heads (h) and tails (t) with six dishes designated for each concentration.

### Drug treatment and Amputation

All concentrations were prepared as a 10mL solution. The ivermectin concentrations used were: control (0μL IVM in 10mL spring water), Pre-soak (0μL IVM in 10mL spring water), 0.5μM (50μL IVM in 9.95mL spring water), 1.0μM (100μL IVM in 9.90mL spring water), 1.0μM (w/o soak) (100μL IVM in 9.90mL spring water), and 5.0μM (500μL IVM in 9.50mL spring water). Planaria treated with 1.0μM (w/o) soak was not exposed to the soak prior to amputation, but received the same concentration of 1μM of IVM treatment. DMSO-treated controls were used to show DMSO not to be a deterrent in regeneration. A Pre-soak consisted of the soak, but no treatment post-amputation. Planaria were transferred into IVM solutions with a colony of 10 in each petri dish right after. Immediately, planaria were bisected horizontally using a sharp scalpel and applying a rocking motion. Planaria were left in the IVM treatments for eight days and observed post amputation for a total of 13 days post soak (Fig. 1).

**Figure 1:**
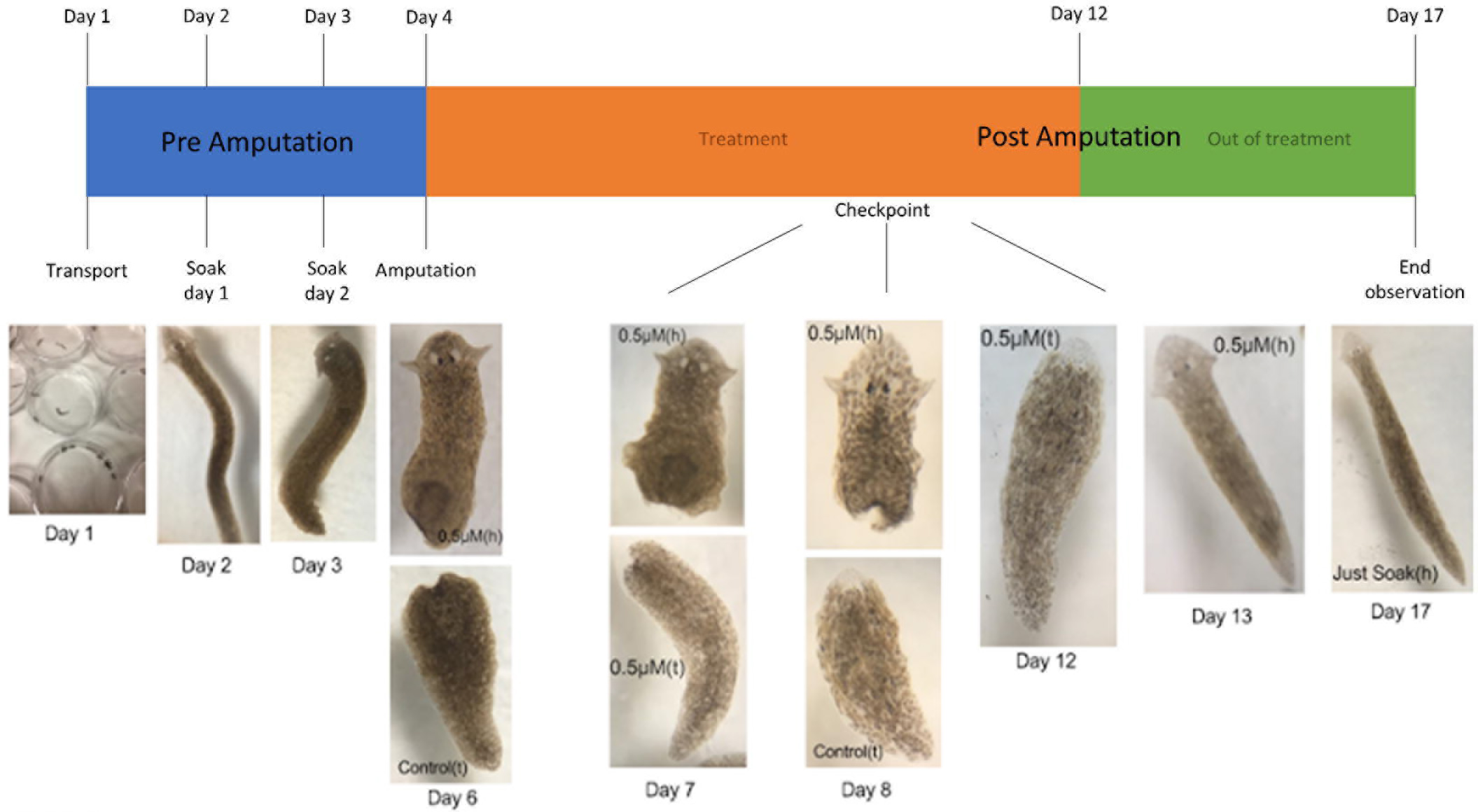
Schematic of the experimental design. The schematic represents the Pre and Post Amputation experiments. Pre Amputation includes transport in which the planaria were transferred from the container they were shipped in into the prepared petri dishes, two day soak, and amputation. Post Amputation includes checkpoints during eight day treatment and observation out of treatment for five days. The total procedure lasted 17 days.

### Image collection and Statistical analysis for phenotypic data

Images were collected using a iPhone camera shot through a Swift M3500D Biological Microscope. T-tests were run through Fathom Dynamic Data Software. Chi-square tests were generated using the Easy Fisher Exact Test Calculator.

## Results

### Establishing a toxicology profile for ivermectin in Dugesia dorotocephala

In order to identify an appropriate (non-toxic) exposure profile for ivermectin in this species of planaria, we examined the mortality incidence over the course of the observation period post amputation of several different concentrations of the drug. These data revealed a correlation between IVM concentration and mortality rates. Planaria treated with IVM for eight days responded with adverse effects of tissue degeneration and death. Higher concentrations of IVM resulted in increased mortality: IVM at 5.0μM was highly toxic and induced fatal reactions within a short timespan (≤48 hours) as well as lysis of planaria (Fig. 2Aii compared to Fig. 2Ai). The fourth day of observation exhibited the highest mortality increase for 1.0μM and 1.0μM (w/o soak) at 41.7%, while the highest mortality increase occurred on ninth day for 0.5μM with 18.3% mortality (Fig. 2B). The control (0μM) and Pre-soak (received a soak, but no treatment post-amputation) resulted in a 0% mortality incidence. Not only did the mortality incidence increase with increasing IVM dosages, but the rate at which these incidences appeared accelerated. However, 0.5μM had the lowest final mortality incidence of 45%, a more suitable concentration to use compared to the other concentrations with mortality incidences of 91.67% (1.0μM), 70% (1.0μM (w/o soak)), and 100% (5.0μM) (Fig. 2C). This established a usable level of IVM exposure that allowed for investigation of subtle patterning phenotypes in the absence of significant general toxicity in subsequent experiments.

**Figure 2:**
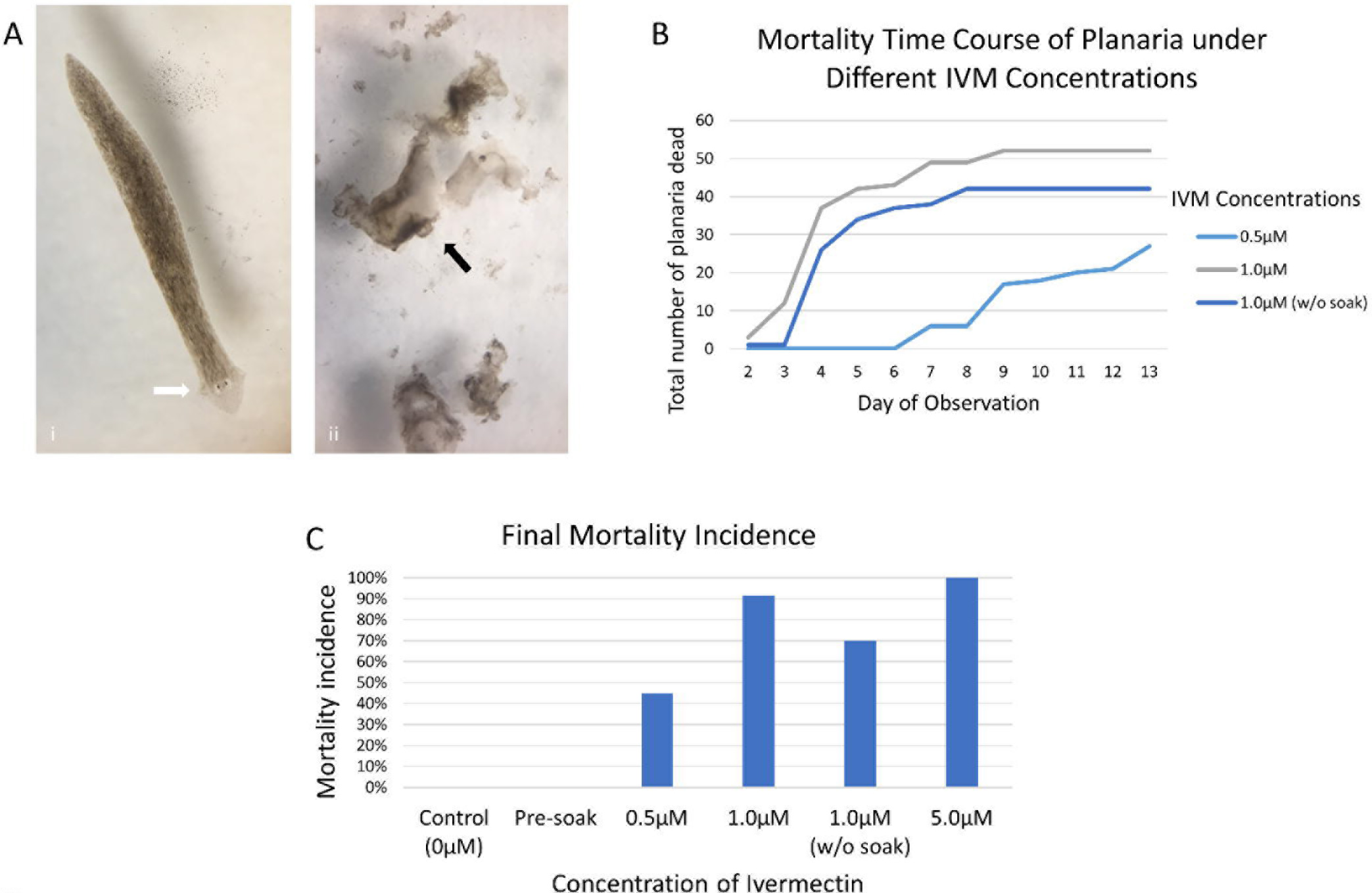
Exposure of ivermectin increases mortality rates in *Dugesia dorotocephala*. *Mortality rate analyses at 13 days regeneration*. **(A)** Representative physical changes are shown. White arrows indicate normal planaria, black represent abnormal, and red represent any missing bodily structure. (panel i) Planaria excluded from IVM exposure (control 0μM) produced no abnormal growth. (panel ii) Lysis of planaria under 5.0μM IVM occurred by the second day post amputation. **(B)** Mortality Time Course of Planaria under Different IVM Concentrations reveal fourth day of post amputation observation to have the highest mortality for 1.0μM and 1.0μM (w/o soak), while the highest mortality occurred on ninth day for 0.5μM. **(C)** Final Mortality Incidence. Control (0μM) (0%), Presoak (0%), 0.5μM (45%), 1.0μM (91.67%), 1.0μM (w/o soak) (70%), and 5.0μM (100%) IVM treated on planaria result in varying mortality rates. N=60 fragments per concentration.

### Ivermectin exposure delays planarian regeneration

In order to determine the regeneration rate of planaria under varying IVM concentrations, we averaged the tracked daily growth of head/tail regeneration. The control (0μM) averaged 8.84 days until full regeneration with a standard deviation of 1.937, Presoak 9.95 with a SD of 1.486, 0.5μM 10.89 with a SD of 1.945. 1.0μM averaged 12 days until full regeneration, 1.0μM (w/o soak) 6, and 5.0μM had no recordable data due to 100% mortality (Fig. 3). Regeneration data was scored using the living planaria. DMSO vehicle was added to the control (0μM) revealing DMSO to not be a deterrent in the regeneration speed. Error bars not shown for experiments where N<5 due to toxicity. The T-test was performed using Fathom Dynamic Data Software on the number of days until full head/tail regeneration of planaria treated with control (0μM), 0.5μM, and 1.0μM IVM; the observed delays in the treated groups were seen to be significant at t<0.0001. We conclude that exposure to IVM delays the regenerative response.

**Figure 3:**
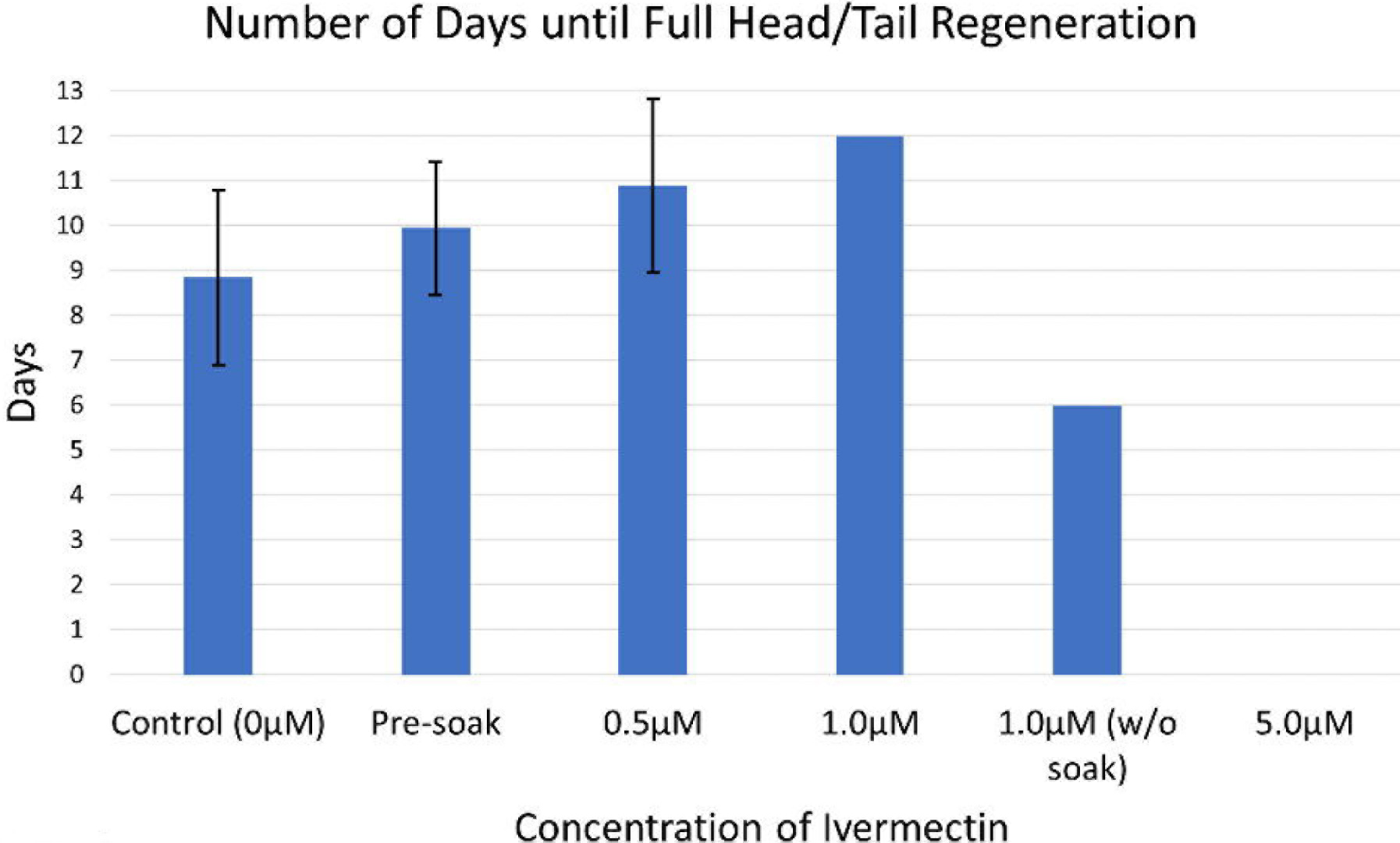
Planaria treated with higher concentrations of IVM exhibit prolonged rates of regeneration. The number of days until full head/tail regeneration was quantified. Under higher concentrations of IVM, the length until full regeneration of planaria increased. Data consisted of only planaria that regenerated. The number of planaria that regenerated for each concentration is as followed: Control (0μM) (59 planaria), Pre-soak (63 planaria; during the experimental set up, more than 30 planaria prior to amputation were included in experimentation for Pre-soak), 0.5μM (21 planaria), 1.0μM (1 planaria), 1.0μM (w/o soak) (1 planaria), 5.0μM (0 planaria). Statistical Analysis by T-test was created using Fathom Dynamic Data Software. T-test indicates significant difference between treated and control groups to p<0.01. N=60 fragments per concentration.

### Ivermectin exposure alters anatomical pattern of planarian regeneration

During the observation period, planaria exposed to IVM deviated from its programed pattern formation and revealed phenotypic abnormalities. Exposure to ivermectin at 1.0μM and 0.5μM induced protrusions along the trunk and anterior of the worms (Fig. 4Aii and iv). Protrusions appeared to be light in color. Planaria regenerating in 0.5μM IVM revealed the greatest variety of altered morphologies, most likely due to decreased toxicity as compared to the higher dosages (Fig. 5A and B). Abnormal morphologies included incomplete regeneration (Fig. 4Av), no regeneration (Fig. 4Aiii), bifurcated tail (Fig. 4Biii), and bifurcated tail with partial head (Fig. 4Bi, ii, and iv). 5.0(μM IVM had no recordable data of morphologies due to the 100% mortality incidence. Phenotypic assessments were made scoring the living planaria. For planaria soaked in treatments of 0.5μM, 1.0μM, and 1.0μM (w/o soak), a chi-square test comparing to controls was significant at p<0.01, thus demonstrating that ivermectin exposure alters morphogenesis during axial patterning in planarian regeneration.

**Figure 4:**
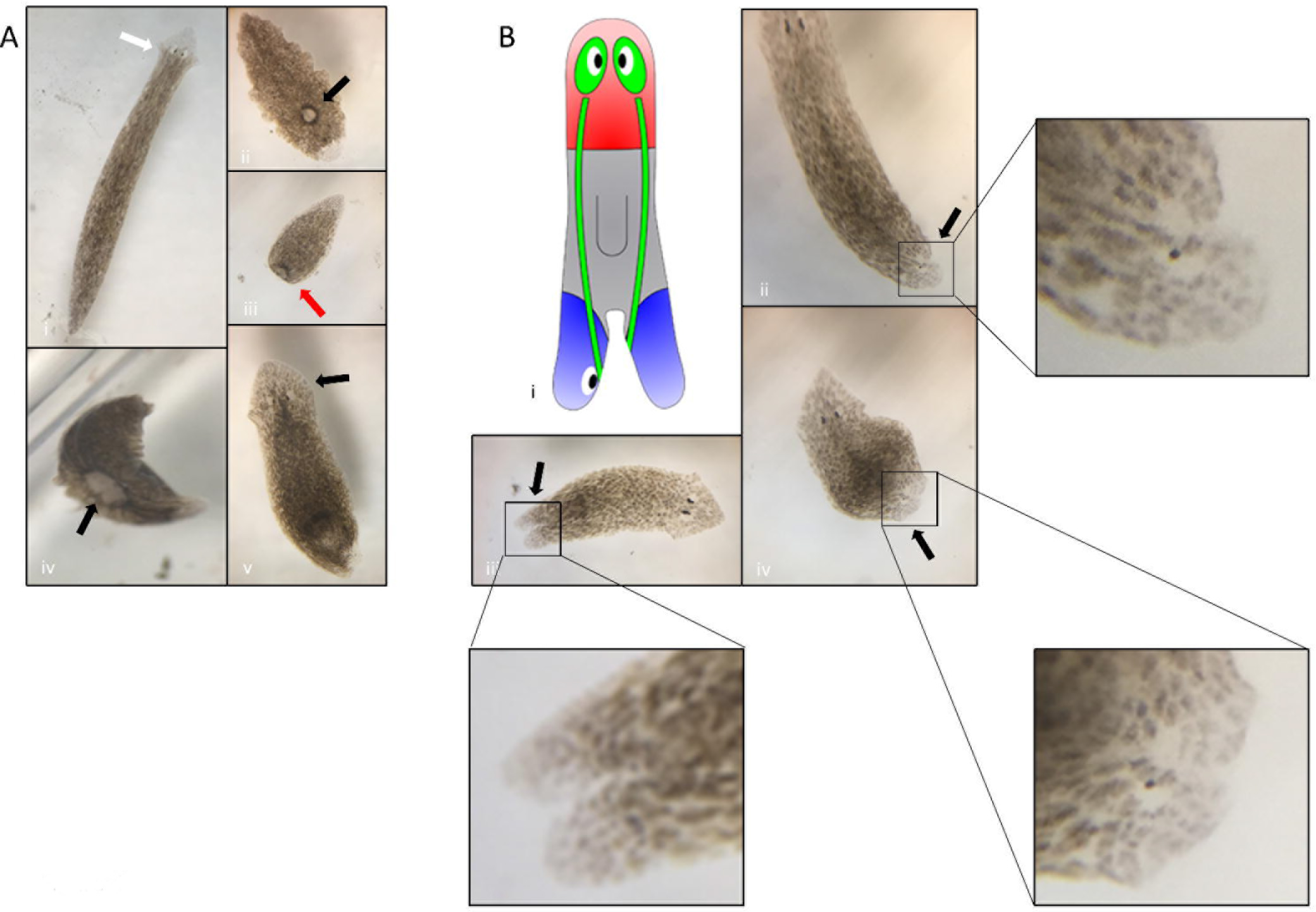
Pattern formation of *Dugesia dorotocephala* during regeneration. **(A)** Planaria abnormal/normal morphology. White arrows represent normal planaria, black represent abnormal, and red represent any missing bodily structure. (panel i) Control (0μM) regenerated normal heads by day 11 post amputation. DMSO was added to the control (0μM) revealing DMSO to not be a deterrent in regeneration. (panel ii) 0.5(μM IVM inhibits regeneration in some planaria. (panel iii and iv) 1.0(μM and 0.5(μM IVM induced protrusions in planaria. The causes are unknown for this abnormality. (panel v) Tail fails to fully regenerate (incomplete regeneration) after full observation in a treatment of 0.5(μM IVM. **(B)** Bifurcated tail in *Dugesia dorotocephala* resulting fro IVM exposure. (panel i) The bifurcated tail with partial head planaria schematic was illustrated schematically using Planform: Planarian Experiment Database and Software Tool - Lobo Lab. (panel ii) 0.5(μM IVM exposure induced bifurcated tail abnormality. (panel iii) 0.5(μM IVM exposure induced bifurcated with partial head on planaria. N=60 fragments per concentration.

**Figure 5:**
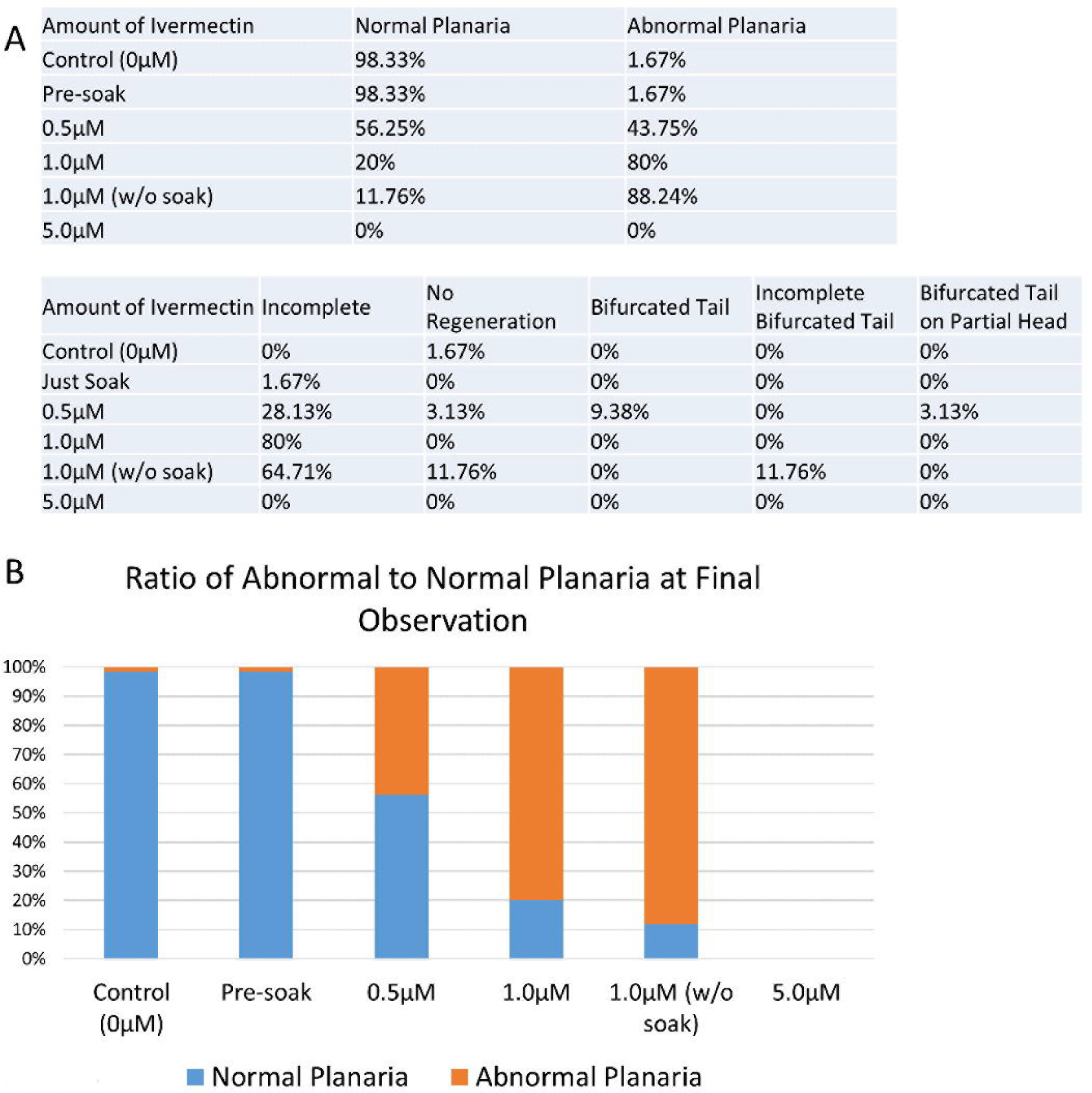
Ivermectin induces abnormalities in *Dugesia dorotocephala*. **(A)** Increasing IVM concentration increases the number of abnormalities. The greatest variety of abnormalities occurred in 0. 5μM IVM, which includes incomplete, no regeneration, bifurcated tail, and bifurcated tail with partial head. 5.0(μM IVM had no recordable data of morphologies due to 100% mortality incidence. **(B)** Ratio of abnormal to normal planaria at the final observation was recorded. The graph illustrates the increasing abnormalities due to the higher concentrations of IVM. N=60 fragments per concentration.

## Discussion

In this study, performed for the first time on *D. dorotocephala,* we demonstrated that exposure to ivermectin, resulted in abnormal pattern formation. Focusing on a new species of planaria, as compared to most other studies using *D. japonica* and *S. mediterranea,* allows for additional novel data on a chloride channel opener drug’s effects. Moreover, establishing a toxicity profile on these planaria facilitates further research on bioelectric properties in this model. As the concentration of IVM increased, abnormalities and mortality increased as well, and the number of days until full head/tail regeneration was delayed. Mortality was especially evident in planaria treated with 5.0μM of IVM. Phenotypic results included protrusions (of unknown tissue composition), partial head, incomplete regeneration, and no regeneration; however, the most noteworthy morphology is the bifurcated tail, reported (to our knowledge) for the first time in this species of planaria. The effects of IVM reveal the role for ivermectin’s target in cells’ patterning decisions during regeneration in this species.

IVM’s alteration of chloride ion flow via voltage membrane was likely responsible for the changes in pattern formation and the protuberances [10], although the existence of additional targets of IVM in this species cannot be ruled out by our data. The most likely mechanism involves a change in flux of chloride ions through the glutamate-gated chloride channels (GluCl), which changed the membrane voltage, a regulator of polarity, initiating partial head formation on the bifurcated tail [26, 45, 46]. Future studies will extend these results by measuring the membrane voltage directly and recording the relative concentration of chloride ions inside vs. outside of the cell. The use of voltage-sensitive reporter dyes, such as DiSBAC2(3), DiSBAC4(3), and CC2-DMPE, as well as expression analysis with molecular markers of regenerative patterning, will enable future characterization of the mechanisms by which changes in channel activity alter pattern regulation during regeneration [2, 39, 40].

Recent experiments have begun to reveal that changes in patterning and morphogenesis are triggered by endogenous bioelectrical changes in ion signaling and bioelectrical gradients [27, 41, 42]. This is due to the fact that patterns of ion flows carry information during planarian regeneration regarding pattern formation; therefore, an altered state of signaling could have induced the changes observed in experimentation [10, 26, 43, 44]. IVM, a chloride channel opener drug, was reported to cause cell depolarization due to membrane voltage disturbances in planaria as well as *C. elegans,* and was shown to be sufficient to alter pattern formation [10, 26, 28].

Anatomical polarity can be altered by bioelectric gradient dynamics,[47–49]. This is due to the spatial patterns of membrane polarization in planaria being important for regeneration; significant alteration can be expected to alter the patterning outcome [26, 44]. In order to understand the establishment of anterior-posterior anatomical polarity and the role of ionic gradients therein, detailed measurements of electrical properties across this species of planaria will be needed, to compare and contrast against available data in the other model species.

Planarians are an excellent model system for identifying new controls of growth and form. It is especially well-suited for the discovery of biophysical signals underlying regeneration, and for the screening of reagents that can be used to manipulate bioelectric signaling. Here, we identified fascinating regenerative phenotypes in a new planarian model species, consistent with a broad role of bioelectric properties in repair and remodeling. A better understanding of how ion channels contribute to patterning decisions will drive important advances in biomedicine addressing traumatic injury, aging, and perhaps other diseases with a significant patterning component (such as cancer and birth defects).

## Acknowledgments

We thank Junji Morokuma, Research Associate for Dr. Michael Levin’s Lab, Allen Discovery Center at Tufts and Tufts Center for Regenerative and Developmental Biology and Department of Biology, for advising us on our amputation procedure; Fayola Lavenhouse, Westfield High School Chemistry teacher at Westfield High School for spending every lunch and Saturday supervising; Rakela Colón, Laboratory Manager at Tufts Center for Regenerative and Developmental Biology for sending us the 10μM of ivermectin in DMSO from Tufts Center for Regenerative and Developmental Biology and Department of Biology; Maya Emmons-Bell Ph.D, Student at UC Berkeley, Beckman Scholar at Tufts University, for advising us on our amputation procedure and for answering our inquiries.

## References

1. Chernet, B.T. and M. Levin, Transmembrane voltage potential is an essential cellular parameter for the detection and control of tumor development in a Xenopus model. Dis Model Mech, 2013. 6(3): p. 595–607.

2. Oviedo, N.J., et al., Live Imaging of Planarian Membrane Potential Using DiBAC4(3).CSH Protoc, 2008. 2008: p. pdb.prot5055.

3. Le, X., et al., Heat shock-inducible Cre/Lox approaches to induce diverse types of tumors and hyperplasia in transgenic zebrafish. Proc Natl Acad Sci U S A, 2007. 104(22): p. 9410–5.

4. Basanta, D., M. Miodownik, and B. Baum, The evolution of robust development and homeostasis in artificial organisms. PLoS Comput Biol, 2008. 4(3): p. e1000030.

5. Birnbaum, K.D. and A. Sanchez Alvarado, Slicing across kingdoms: regeneration in plants and animals. Cell, 2008. 132(4): p. 697–710.

6. Karami, A., et al., Planarians: an In Vivo Model for Regenerative Medicine. Int J Stem Cells, 2015. 8(2): p. 128–33.

7. Baguna, J., The planarian neoblast: the rambling history of its origin and some current black boxes. Int J Dev Biol, 2012. 56(1-3): p. 19–37.

8. Rink, J.C., Stem cell systems and regeneration in planaria. Dev Genes Evol, 2013. 223(1-2): p. 67–84.

9. Sheiman, I.M. and I.D. Kreshchenko, [Regeneration of planarians: experimental object]. Ontogenez, 2015. 46(1): p. 312.

10. Lobo, D., W.S. Beane, and M. Levin, Modeling planarian regeneration: a primer for reverse-engineering the worm. PLoS Comput Biol, 2012. 8(4): p. e1002481.

11. Salo, E., et al., Planarian regeneration: achievements and future directions after 20 years of research. Int J Dev Biol, 2009. 53(8-10): p. 1317–27.

12. Levin, M., Large-scale biophysics: ion flows and regeneration. Trends Cell Biol, 2007. 17(6): p. 261–70.

13. Sullivan, K.G., M. Emmons-Bell, and M. Levin, Physiological inputs regulate species-specific anatomy during embryogenesis and regeneration. Commun Integr Biol, 2016. 9(4): p. e1192733.

14. Levin, M., G. Pezzulo, and J.M. Finkelstein, Endogenous Bioelectric Signaling Networks: Exploiting Voltage Gradients for Control of Growth and Form. Annu Rev Biomed Eng, 2017. 19: p. 353–387.

15. Ikeda, T., Pharmacological effects of ivermectin, an antiparasitic agent for intestinal strongyloidiasis: its mode of action and clinical efficacy. Nihon Yakurigaku Zasshi, 2003. 122(6): p. 527–38.

16. Blackiston, D., et al., Transmembrane potential of GlyCl-expressing instructor cells induces a neoplastic-like conversion of melanocytes via a serotonergic pathway. Dis Model Mech, 2011. 4(1): p. 67–85.

17. Blackiston, D.J., et al., A novel method for inducing nerve growth via modulation of host resting potential: gap junction-mediated and serotonergic signaling mechanisms. Neurotherapeutics, 2015. 12(1): p. 170–84.

18. Pai, V.P., et al., Endogenous gradients of resting potential instructively pattern embryonic neural tissue via Notch signaling and regulation of proliferation. J Neurosci, 2015. 35(10): p. 4366–85.

19. Anderson, J.D., et al., Vcompoundoltage-gated sodium channel blockers as cytostatic inhibitors of the androgen-independent prostate cancer cell line PC-3. Mol Cancer Ther, 2003. 2(11): p. 1149–54.

20. Sikes, R.A., et al., Therapeutic approaches targeting prostate cancer progression using novel voltage-gated ion channel blockers. Clin Prostate Cancer, 2003. 2(3): p. 181–7.

21. Wang, X.T., et al., The mRNA of L-type calcium channel elevated in colon cancer: protein distribution in normal and cancerous colon. Am J Pathol, 2000. 157(5): p. 1549–62.

22. Preussat, K., et al., Expression of voltage-gated potassium channels Kv1.3 and Kv1.5 in human gliomas. Neurosci Lett, 2003. 346(1-2): p. 33–6.

23. Laniado, M.E., S.P. Fraser, and M.B. Djamgoz, Voltage-gated K(+) channel activity in human prostate cancer cell lines of markedly different metastatic potential: distinguishing characteristics of PC-3 and LNCaP cells. Prostate, 2001. 46(4): p. 262–74.

24. Schonherr, R., Clinical relevance of ion channels for diagnosis and therapy of cancer. J Membr Biol, 2005. 205(3): p. 175–84.

25. Fiske, J.L., et al., Voltage-sensitive ion channels and cancer. Cancer Metastasis Rev, 2006. 25(3): p. 493–500.

26. Beane, W.S., et al., A chemical genetics approach reveals H,K-ATPase-mediated membrane voltage is required for planarian head regeneration. Chem Biol, 2011. 18(1): p. 77–89.

27. Durant, F., et al., Long-Term, Stochastic Editing of Regenerative Anatomy via Targeting Endogenous Bioelectric Gradients. Biophys J, 2017. 112(10): p. 2231–2243.

28. Pemberton, D.J., et al., Characterization of glutamate-gated chloride channels in the pharynx of wild-type and mutant Caenorhabditis elegans delineates the role of the subunit GluCl-alpha2 in the function of the native receptor. Mol Pharmacol, 2001. 59(5): p. 1037–43.

29. Tuck, E. and V. Cavalli, Roles of membrane trafficking in nerve repair and regeneration. Vol. 3. 2010. 209–14.

30. O’Donovan, K.J., Intrinsic Axonal Growth and the Drive for Regeneration. Frontiers in Neuroscience, 2016. 10(486).

31. Levin, M., Gap junctional communication in morphogenesis. Progress in Biophysics and Molecular Biology, 2007. 94(1): p. 186–206.

32. Barghouth, P.G., M. Thiruvalluvan, and N.J. Oviedo, Bioelectrical regulation of cell cycle and the planarian model system. Biochimica et Biophysica Acta (BBA) - Biomembranes, 2015. 1848(10): p. 2629–2637.

33. Levin, M., Molecular bioelectricity in developmental biology: new tools and recent discoveries: control of cell behavior and pattern formation by transmembrane potential gradients. Bioessays, 2012. 34(3): p. 205–17.

34. Oviedo, N.J., et al., Establishing and maintaining a colony of planarians. CSH Protoc, 2008. 2008: p. pdb.prot5053.

35. Alonso, A. and J.A. Camargo, Ameliorating effect of chloride on nitrite toxicity to freshwater invertebrates with different physiology: a comparative study between amphipods and planarians. Arch Environ Contam Toxicol, 2008. 54(2): p. 259–65.

36. Pagan, O.R., T. Coudron, and T. Kaneria, The flatworm planaria as a toxicology and behavioral pharmacology animal model in undergraduate research experiences. J Undergrad Neurosci Educ, 2009. 7(2): p. A48–52.

37. Pagan, O.R., A.L. Rowlands, and K.R. Urban, Toxicity and behavioral effects of dimethylsulfoxide in planaria. Neurosci Lett, 2006. 407(3): p. 274–8.

38. Paskin, T.R., et al., Planarian Phototactic Assay Reveals Differential Behavioral Responses Based on Wavelength. PLoS One, 2014. 9(12): p. e114708.

39. Adams, D.S. and M. Levin, Measuring resting membrane potential using the fluorescent voltage reporters DiBAC4(3) and CC2-DMPE. Cold Spring Harb Protoc, 2012. 2012(4): p. 459–64.

40. Adams, D.S. and M. Levin, General principles for measuring resting membrane potential and ion concentration using fluorescent bioelectricity reporters. Cold Spring Harb Protoc, 2012. 2012(4): p. 385–97.

41. Levin, M., Bioelectric mechanisms in regeneration: Unique aspects and future perspectives. Semin Cell Dev Biol, 2009. 20(5): p. 543–56.

42. Beane, W.S., et al., Bioelectric signaling regulates head and organ size during planarian regeneration. Development, 2013. 140(2): p. 313–22.

43. Nogi, T., et al., Eye regeneration assay reveals an invariant functional left-right asymmetry in the early bilaterian, Dugesia japonica. Laterality, 2005. 10(3): p. 193–205.

44. Nogi, T., et al., A novel biological activity of praziquantel requiring voltage-operated Ca2+ channel beta subunits: subversion of flatworm regenerative polarity. PLoS Negl Trop Dis, 2009. 3(6): p. e464.

45. Shan, Q., J.L. Haddrill, and J.W. Lynch, Ivermectin, an unconventional agonist of the glycine receptor chloride channel. J Biol Chem, 2001. 276(16): p. 12556–64.

46. Schatzberg, D., et al., H(+)/K(+) ATPase activity is required for biomineralization in sea urchin embryos. Dev Biol, 2015. 406(2): p. 259–70.

47. Dimmitt, J. and G. Marsh, Electrical control of morphogenesis in regenerating Dugesia tigrina. II. Potential gradient vs. current density as control factors. J Cell Comp Physiol, 1952. 40(1): p. 11–23.

48. Adell, T., F. Cebria, and E. Salo, Gradients in planarian regeneration and homeostasis. Cold Spring Harb Perspect Biol, 2010. 2(1): p. a000505.

49. Marsh, G. and H.W. Beams, Electrical control of morphogenesis in regenerating Dugesia tigrina. I. Relation of axial polarity to field strength. J Cell Comp Physiol, 1952. 39(2): p. 191–213.

